# ATP2B1 expression identifies human hematopoietic stem cells across ontogeny with superior repopulation and self-renewal capacity

**DOI:** 10.1101/2025.06.11.659017

**Authors:** Angelica Varesi, Murtaza S. Nagree, Isabella Di Biasio, Andy G.X. Zeng, Sayyam Shah, Hyerin Kim, Michael Zhang, Alex Murison, John E. Dick, Stephanie Z. Xie

## Abstract

Long-term hematopoietic stem cells (LT-HSC) maintain lifelong hematopoiesis while preserving the stem cell compartment through self-renewal. The human LT-HSC compartment is molecularly and functionally heterogeneous and also varies across ontogeny. Dissecting the molecular basis for this variation is impeded by the absence of immunophenotypic markers to resolve LT-HSC heterogeneity. Here, we identified ATPase plasma membrane Ca^2+^transporting 1 (ATP2B1/PMCA1) as a novel cell surface marker that is heterogeneously expressed by CD49f^+^ LT-HSC across ontogeny. ATP2B1 immunophenotypic expression stratified human CD49f^+^ LT-HSC from fetal liver (FL), neonatal cord blood (CB) and adult mobilized peripheral blood (mPB) sources into functionally distinct subpopulations in single-cell (sc) clonogenic assays. CD49f^+^ATP2B1^+^ LT-HSC exhibited superior long-term repopulation and self-renewal capacities *in vivo* compared to CD49f^+^ATP2B1^–^ LT-HSC. Molecular profiling by scMultiome and immunofluorescence microscopy point to enrichment of an HSC self-renewal program that includes the TFEB-endolysosomal axis in CD49f^+^ATP2B1^+^ LT-HSC. Our study provides a new tool to dissect the heterogeneous molecular programs in LT-HSC across ontogeny.

## Introduction

Human hematopoiesis is highly dynamic, originating from a rare pool of dormant long-term hematopoietic stem cells (LT-HSC) that both self-renew and generate downstream progeny.^1,2^ In contrast to the mouse system, immunophenotypic markers that permit isolation of human LT-HSC to single unit purity have not been reported. Beginning with isolation of CD34 expressing cells, a number of HSC-sorting schemes have been developed. Expression of CD49f (integrin α6) and CD90 is most commonly used to resolve hematopoietic stem-enriched cells (CD34^+^CD38^−^CD45RA^−^) into short-term HSC (ST-HSC, CD90^−^CD49f^-^) and LT-HSC (CD90^+^CD49f^+^).^3–5^ Subsequent studies have tried to resolve CD90^+^CD49f^+^ LT-HSC with the use of additional markers such as EPCR,^6^ GPRC5C,^7^ CD112,^8^ and CLEC9A.^9^

These studies have contributed fundamental understanding to the molecular underpinnings of human LT-HSC biology. Nonetheless, functional analyses demonstrated that all these markers still capture a heterogeneous population of LT-HSC with enriched yet variable stem cell frequency. Additionally, most studies are primarily performed using cord-blood (CB) as the HSC source.^8–11^ Comprehensive studies into LT-HSC functional heterogeneity across the human lifespan remain incomplete. Developmental stage studies have shown that fetal HSC are both more proliferative and self-renewing than neonatal CB HSC, while adult HSC are comparatively more quiescent and exhibit less robust stemness properties.^12–19^ Thus, a common marker set that enables prospective separation of functionally distinct human CD90^+^ HSC subsets across ontogeny is needed.

Functional interrogation into the demand-adapted regulatory circuits of the early stages of human hematopoiesis have revealed extensive transcriptional, epigenetic, and metabolic heterogeneity in human HSC.^4,7–10,19–25^ Specifically, studies in CB uncovered catabolic programs centred around TFEB-mediated control of endolysosomal activity as crucial for quiescence and self-renewal in LT-HSC.^10^ Lysosomal activity was shown to distinguish self-renewal potential of LT-HSC.^10^ Yet, whether such programs are specific only to CB-derived LT-HSC or are conserved throughout human development from FL to adult LT-HSC is unknown. Moreover, a surface marker distinguishing LT-HSC with active TFEB at homeostasis is lacking. Lysosomes also act as a major intracellular signaling hub, in part functioning as a major intracellular store of Ca^2+^.^26^ Interestingly, murine and human HSC contain lower intracellular Ca^2+^ levels compared to committed progenitors, and *ex vivo* culture in low Ca^2+^ conditions preserves stemness.^27^ The Ca^2+^exporter ATPase plasma membrane Ca^2+^ transporting 1 (*ATP2B1/PMCA1*) has emerged as a key regulator of mouse embryonic stem cell pluripotency,^26,28^ but its relevance in human stem cells remains unexplored.

Here, we refine the characterization of the human hematopoietic hierarchy by identifying ATP2B1 as a novel immunophenotypic marker that stratifies human CD49f^+^ LT-HSC cells into two distinct functional compartments across ontogeny. Through a comprehensive set of single-cell *in vitro* assays, single-cell Multiomic (scMultiome) molecular analysis, and *in vivo* xenograft functional studies, we show that CD49f^+^ATP2B1^+^ LT-HSC have sustained long-term repopulation capacity and superior self-renewal ability with balanced multilineage output.

## Results

### ATP2B1 is expressed on the surface of human LT-HSC across ontogeny

To discover new surface proteins able to prospectively isolate functionally distinct subsets of human CD90^+^ HSC across ontogeny, we analyzed various joint single-cell (sc) transcriptional and epigenetic (scMultiome) data sets.^25,29^ As quiescence and dormancy are molecular features associated with LT-HSC function,^10,21^ we restricted our analysis to data sets enriched for primitive populations, and we pinpointed ATP2B1 as a potential surface marker of human LT-HSC. *ATP2B1* expression was significantly higher in HSC enriched for a quiescence vs activation signature after recovery from inflammation in xenografts (Figure S1A).^25^ Further analysis of CB HSPC populations profiled by bulk RNAseq revealed increased ATP2B1 expression in quiescent HSPC compared to activated HSPC fractions (Figure S1B).

To determine if ATP2B1 is expressed at the protein level in human HSC at homeostasis,, we analyzed CB LT-HSC (CD19^−^CD34^+^CD38^−^CD45RA^−^CD90^+^CD49f^+^) and ST-HSC (CD90^−^CD49f^−^) by immunofluorescence microscopy (Figure 1A). ATP2B1 surface expression was variable across cells but covered a larger surface area in LT-HSC compared to ST-HSC (Figure 1B). Next, we analyzed the surface expression of ATP2B1 across the HSPC hierarchy using another ATP2B1 antibody by flow cytometry analysis in CD34^+^ HSPC from FL, CB, and mPB donors (age 22-58 years) to assess if ATP2B1 heterogeneous expression extended beyond CB sources (Figure S1C). The CD90^+^ HSC compartment was partitioned into 3 subpopulations based on the expression of CD49f and ATP2B1: CD49f^+^ATP2B1^+^, CD49f^+^ATP2B1^−^ and CD49f^−^ATP2B1^−^ compared to fluorescence minus one (FMO) controls lacking ATP2B1 antibody (Figures 1C and S1D). The percentage of CD49f^+^ATP2B1^+^ cells decreases across ontogeny with no change from young adults to aged donors (Figure 1D); yet ATP2B1 expression was higher on LT-HSC compared to ST-HSC (Figure 1E) and other downstream committed populations across all three cell sources (Figure S1E,F). Overall, these results show that human CD49f^+^ LT-HSC can be further subdivided into different populations based on ATP2B1 expression.

**Figure 1:**
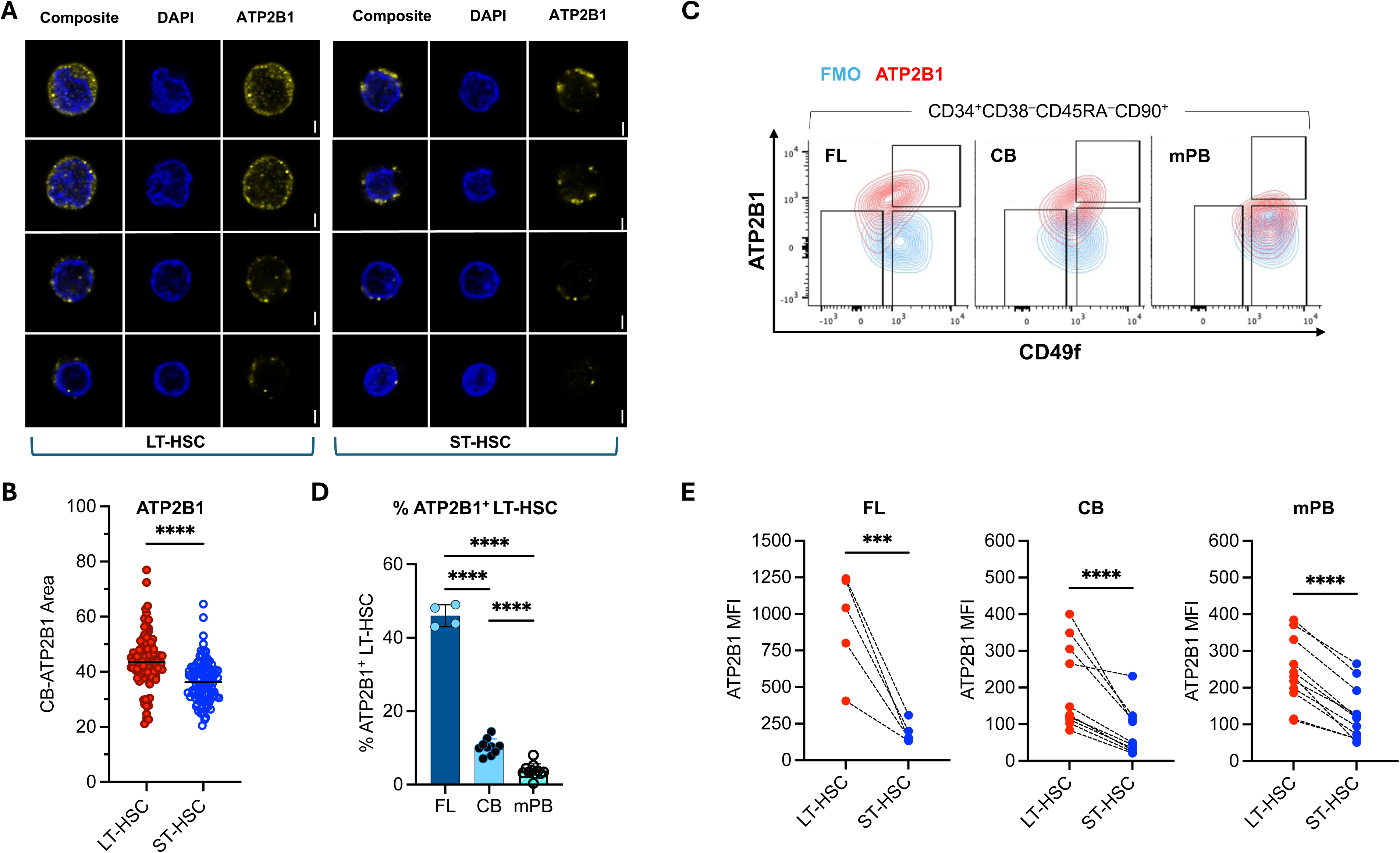
ATP2B1 is expressed on the surface of human LT-HSC across ontogeny. (**A,B**): Representative images (**A**) and quantification (**B**) of confocal analysis of LT-HSC and ST-HSC stained for ATP2B1 and DAPI. Scale bar, 2 μm. n = 2 CB,195 individual cells/staining. Mann-Whitney test. Scatter dot plot. Median with interquartile range. : Representative flow cytometric plots of ATP2B1^+/–^ and CD49f^+/–^ cells within human CD34^+^CD38^−^CD45RA^−^CD90^+^ HSC in FL, CB and mPB. Light blue: FMO: Fluorescent Minus One; red: ATP2B1 stained cells. : Percentage of CD49f^+^ATP2B1^+^cells. n=4 FL, n=10 CB, n=10 mPB. One Way Anova. Scatter dot plot. Mean with SD. : ATP2B1 MFI in LT-HSC and ST-HSC analyzed after thawing by flow cytometry. n=4 FL, n=10 CB, n=10 mPB. Ratio paired t test. Before-after. Symbols & lines.

### ATP2B1^+^ LT-HSC have superior *in vitro* clonogenic potential

To investigate functional differences among the newly defined HSC subsets across ontogeny, Fluorescence-Activated Cell Sorting (FACS) was used to isolate CD49f^+^ATP2B1^+^, CD49f^+^ATP2B1^−^ and CD49f^−^ATP2B1^−^ HSC from FL, CB and mPB (n=5 FL, 14 CB, 10 mPB donors) followed by serial *in vitro* clonogenic assays (Figure S2A). At day 14, the total number of colonies was the highest in FL (mean 47.6 colonies) compared to CB (mean 29.5 colonies, *p*<0.0087) and mPB (mean 7 colonies, *p*<0.0001) samples, in line with previous reports of superior clonogenic output in FL CD49f^+^ LT-HSC compared to CB and mPB with single-cell stromal differentiation assays (Figure S2B).^19^ For both CB and mPB, the clonogenic potential of the CD49f^+^ATP2B1^+^ subset was significantly lower than the CD49f^+^ATP2B1^−^ and CD49f^−^ATP2B1^−^ subsets (Figure 2A). No significant differences in clonogenic output were observed between the 3 populations in FL (Figure 2A). However, flow cytometry analysis showed the total proportion of primitive CD34^+^ cells in the primary colony assay was highest in the CD49f^+^ATP2B1^+^ subset from all HSC sources and then decreased from CD49f^+^ATP2B1^−^ to CD49f^−^ATP2B1^−^ (Figure 2B). Importantly, serial replating assays resulted in a higher clonogenic output from CD49f^+^ATP2B1^+^ compared to CD49f^+^ATP2B1^−^ and CD49f^−^ATP2B1^−^ progeny for all three human HSC sources (Figure 2C). Despite some differences in erythroid, granulocyte, or myeloid lineage output among the conditions tested, they were mostly lost upon serial repeating and not consistent across ontogeny (Figure S2C,D). These data suggest that CD49f^+^ATP2B1^+^ cells contain the highest long-term clonogenic potential in the HSC compartment without impairment in their multilineage clonogenic capacity.

**Figure 2:**
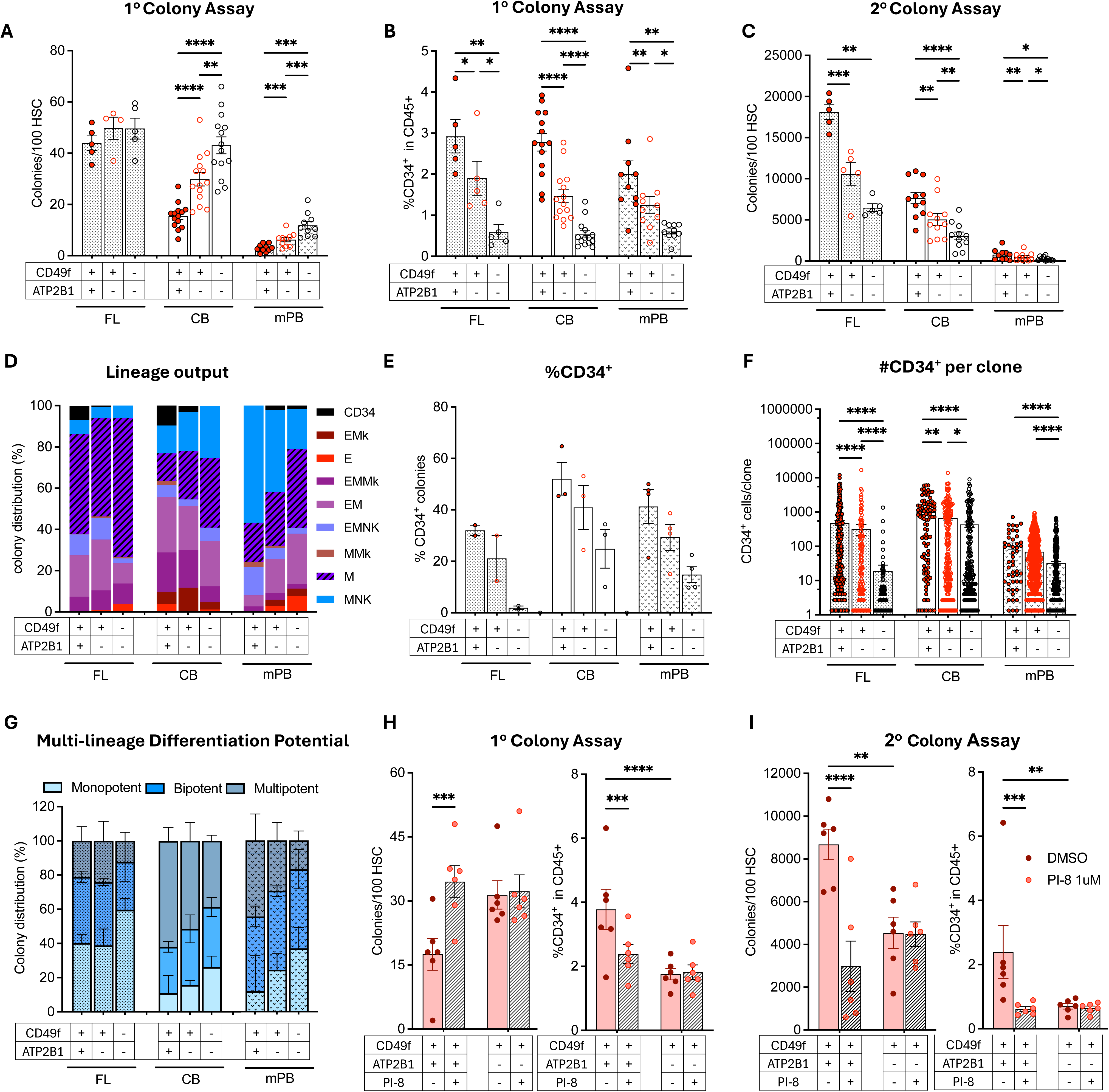
ATP2B1^+^ LT-HSC have superior *in vitro* clonogenic potential . (**A-C**): CD49f^+^ATP2B1^+^, CD49f^+^ATP2B1^−^ and CD49f^−^ATP2B1^−^ clonogenic assays showing (**A**) number of primary colonies/100 cells (n=5 FL, 14 CB, 10 mPB); (**B**) number of secondary colonies/100 cells (n=5 FL, 14 CB, 10 mPB); (**C**) percentage of CD34^+^ within CD45^+^ cells from primary colonies in A. One Way Anova test. p>0.05 is not shown. Scatter dot plot. Mean with SEM. (**D**): Lineage output from the in vitro single-cell differentiation assay of individual HSCfrom FL (n=2), CB (n=3) and mPB (n=4) experiments with independent single donors (FL and mPB) or donor pools (CB). Numbers of single-cell colonies with positive genotype are indicated at each condition (CD34^+^: CD34^+^ only colonies, E = erythroid (CD45^−^/GlyA^+^), M = myeloid (CD14^+^ and/or CD66^+^ and/or CD33^+^), Mk = megakaryocytic (CD41^+^), NK: CD56^+^ cells. (**E**): Percentage of colonies containing CD34^+^ cells in the single-cell in vitro differentiation assays from FL (n=2), CB (n=3) and mPB (n=4) experiments with independent single donors (FL and mPB) or donor pools (CB). Scatter dot plot. Mean with SEM. (**F**): Number of CD34^+^ cells per clone in the single-cell in vitro differentiation assays from FL (n=2), CB (n=3) and mPB (n=4) experiments with independent single donors (FL and mPB) or donor pools (CB). Kruskal Wallis test. p>0.05 is not shown. Box and whiskers. Min to max show all points. (**G**): Percentage of single cells that grew into single colonies of one lineage (Monopotent), two (Bipotent), and three or more (Multipotent) lineages in the single-cell in vitro differentiation assays from FL (n=2), CB (n=3) and mPB (n=4) experiments with independent single donors (FL and mPB) or donor pools (CB). Bar. Mean with SD. (**H,I**): CD49f^+^ATP2B1^+^ and CD49f^+^ATP2B1^−^ clonogenic assays showing (**H left**) primary and (**I left**) secondary colonies/100 cells in presence of Ctrl or 1 μM PI-8 (n=6, CB) or (**H-I right**) percentage of CD34^+^ within CD45^+^ cells from primary colonies in (**H left**) and secondary colonies in (**I left**). One Way Anova test. p>0.05 is not shown. Scatter dot plot. Mean with SEM.

To assess how ATP2B1 surface expression impacted lineage output at single-cell resolution, single-cell stroma-based differentiation assays were performed from index-sorted FL (n=1280), CB (n=1858) and mPB (n=5036) CD90^+^ HSC (Figure S2E). The cloning efficiency from these stromal-based assays was higher in FL (mean 88.1%) than CB (mean 58.59%) and mPB (mean 41.2%) (Figure S2F). Within those, CD49f^−^ATP2B1^−^ had the highest clonogenicity followed by CD49f^+^ATP2B1^−^ then CD49f^+^ATP2B1^+^ HSC in CB and mPB, but not FL, in line with results from the primary colony-forming assays (Figure S2F). Broadly, the lineage output between the colonies derived from each of the three HSC subpopulations was comparable (Figures 2D and SG-I). Instead, the incidence of clones containing exclusively CD34^+^ cells followed an interesting pattern. In FL and CB, but not in mPB, the abundance of these clones was higher from HSC classified as CD49f^+^ATP2B1^+^ compared to a reduced number from HSC classified as CD49f^+^ATP2B1^−^; they were absent in colonies derived from CD49f^−^ATP2B1^−^ HSC (Figure 2E). We therefore sought to investigate CD34^+^ progeny in more detail. The percentage of CD34^+^ cells was the highest in clones derived from CD49f^+^ATP2B1^+^ than CD49f^+^ATP2B1^−^and CD49f^−^ATP2B1^−^ HSC across all 3 sources (Figure 2E). Moreover, the number of CD34^+^ cells per clone was the highest in CD49f^+^ATP2B1^+^ and gradually decreased in CD49f^+^ATP2B1^−^ and CD49f^−^ATP2B1^−^ HSC (Figure 2F), despite observing no difference in total number of cells per clone between them (Figure S2J). We did not find any clear correlation between ATP2B1 mean fluorescence intensity in the parent cell and the number of CD34^+^ progeny, suggesting the absolute level of expression of ATP2B1 does not contribute to this phenotype (Figure S2K). Importantly, ATP2B1^+^LT-HSC gave rise to 21% (FL), 43% (CB) and 44% (mPB) multi-lineage colonies, with three or more of the following lineages: CD34^+^, erythroid, myeloid, megakaryocytes, and natural killer (Figure2G). This was consistent across biological replicates and lineages, but particularly marked in CB and mPB. (Figure S2L-N).

To determine whether ATP2B1 Ca^2+^ transport activity is functionally important for the higher long-term clonogenic potential of the CD49f^+^ATP2B1^+^ subset, we focused on CB-derived CD49f^+^ LT-HSC and performed colony forming assays in the presence or absence of an ATP2B1 inhibitor, PI-8.^30^ PI-8 treatment significantly increased colony output from the CD49f^+^ATP2B1^+^ in the first plating, and decreased output in secondary plating, phenocopying colony output of the CD49f^+^ATP2B1^−^ population (Figure 2H,I). In agreement, the percentage of CD34^+^ cells in both primary and secondary colonies from CD49f^+^ATP2B1^+^ samples following PI-8 treatment was decreased compared to control (Figure 2H,I). Instead, both clonogenicity and CD34^+^ abundance were not impacted by PI-8 treatment of CD49f^+^ATP2B1^−^ HSC (Figure 2H,I). No significant skewing in lineage output was observed between treated and non-treated colonies (Figure S2O,P). These results suggest ATP2B1 activity has a functional role in the superior clonogenic output demonstrated by CD49f^+^ATP2B1^+^ LT-HSC *in vitro*.

Overall, these data show that ATP2B1 surface expression identifies a subset of ATP2B1^+^CD49f^+^ LT-HSC with superior long-term clonogenic capacity and that the transporter activity of ATP2B1 is required for the distinct functional differences between ATP2B1^+^CD49f^+^ and CD49f^+^ATP2B1^−^ LT-HSC .

### Transcriptional and epigenetic features of CD49f^+^ATP2B1^+^ LT-HSC are enriched for TFEB-regulated lysosomal programs

To gain insight into molecular programs governing the superior long-term clonogenic potential of CD49f^+^ATP2B1^+^ HSC, we performed scMultiome profiling of FACS-purified CD49f^+^ATP2B1^+^, CD49f^+^ATP2B1^−^ and CD49f^−^ATP2B1^−^ CB-HSC using the 10x Multiome platform (Figure S3A). As we observed delayed clonogenic output *in vitro* assays, we hypothesized that ATP2B1 would demarcate more quiescent HSC, a corollary of superior HSC stemness.^4,8,10,21^ We have previously defined functionally validated transcriptional and epigenetic signatures that distinguish quiescent LT-HSPC from activated (Act)-HSPC.^10,21^ We computed epigenetic and transcriptional enrichment scores for these signatures to assess molecular facets of quiescence in our data.^31,32^ Epigenetic features marking LT-HSPC were enriched in CD49f^+^ATP2B1^+^ compared to the other HSC subsets (Figures 3A and S3B). Moreover, the qLTvsST transcriptional signature was concurrently enriched in CD49f^+^ATP2B1^+^ HSC, indicating they are in a more quiescent state than CD49f^+^ATP2B1^−^ and CD49f^−^ATP2B1^−^ HSC (Figure3B). Transcriptional programs were extracted using linked non-negative matrix factorization (L-NMF).^33^ We identified 23 signatures, 20 of which were variably expressed across CD49f^+^ATP2B1^+^, CD49f^+^ATP2B1^−^ and CD49f^−^ATP2B1^−^ HSC. Three signatures showed significant differential enrichment across the HSC subsets: L-NMF16 was enriched in CD49f^+^ATP2B1^+^ HSC, L-NMF6 had increasing expression from CD49f^+^ATP2B1^+^ to CD49f^−^ATP2B1^−^ HSC, and L-NMF23, which was highly enriched in CD49f^−^ATP2B1^−^, was absent in CD49f^+^ATP2B1^+^ and was expressed at a lower level in CD49f^+^ATP2B1^−^ HSC (Figures 3C-E and S3C). Gene set enrichment analysis (GSEA) revealed that the top driving genes within L-NMF16 were enriched in autophagy, catabolism, endolysosomal trafficking and lysosome-related programs (Figure 3F). Conversely, gene sets marking cellular proliferation and stem cell differentiation were mostly associated with L-NMF6 (Figure S3D). To understand the epigenetic traits linked to L-NMF16 and L-NMF6, their enrichment in single cells was correlated to the LT/HSPC and Act/HSPC chromatin signature enrichment in those cells.^21^ LT/HSPC chromatin signatures were enriched in L-NMF16-expressing cells, whereas the Act/HSPC chromatin signature was enriched in L-NMF6 expressing cells (Figure S3E). We then focused on the chromatin differences between CD49f^+^ATP2B1^+^ and CD49f^+^ATP2B1^−^ HSC to gain more granularity within the LT/HSPC pool. Thus, scATAC peaks enriched in CD49f^+^ATP2B1^+^ and CD49f^+^ATP2B1^−^ HSC were intersected with bedtools to obtain unique chromatin accessibility sites in these two LT-HSC subsets. CD49f^+^ATP2B1^+^ cells had 10800 unique sites; 9450 sites were exclusively found in CD49f^+^ATP2B1^−^ HSC; and 43916 sites were common to both. Analysis of transcription factor (TF) recognition motif and accessibility revealed CTCF and BORIS binding sites as most significantly enriched in CD49f^+^ATP2B1^−^ cells,^21^ whereas there was no enrichment of TF motifs over the sites unique toCD49f^+^ATP2B1^+^ HSC (Figure S3F). CD49f^+^ATP2B1^−^ HSC also showed enrichment of characteristics of TF activity associated with anabolic cells, such as that of MYC(Figure S3F).^10^ Conversely, CD49f^+^ATP2B1^+^ and CD49f^+^ATP2B1^−^ LT-HSC showed similar accessibility of HOXB7, a TF common to both LT- and ST-HSC (Figure S3F).^10^ Gene regulatory domains falling within sites unique to CD49f^+^ATP2B1^+^ HSC revealed enrichment in gene ontology (GO) pathways of catabolism, endosomal vesicle trafficking, and autophagy (Figure 3G). Conversely, sites unique to CD49f^+^ATP2B1^−^ were enriched in mitogenic and proliferative pathways which are typical hallmarks of differentiating stem cells (Figure S3G).^10^ Thus, through two independent transcriptional and chromatin accessibility analyses we have uncovered distinct transcriptional and epigenetic programs in ATP2B1^+^ and ATP2B1^−^ LT-HSC that are linked to stemness and these include endolysosome-related pathways.

**Figure 3:**
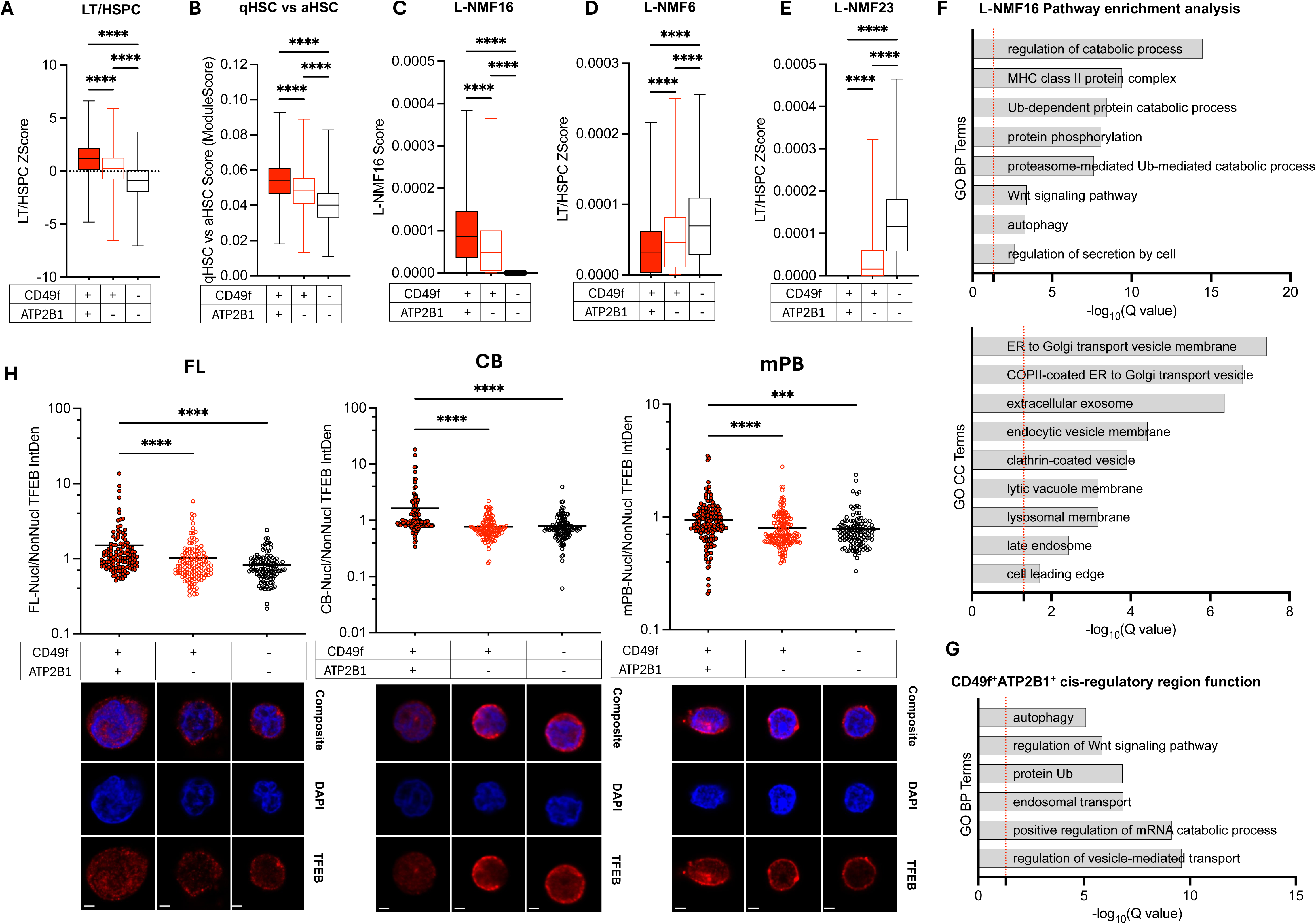
Transcriptional and epigenetic features of CD49f^+^ATP2B1^+^ LT-HSC are enriched for TFEB-regulated lysosomal programs. (**A-E**): Distribution of (**A**) *Z* scores enrichment of the LT/HSPC epigenetic signature, (**B**) ModuleScore enrichment of the qHSC versus aHSC transcriptional signature, (**C**) L-NMF16 enrichment score, (**D**) L-NMF6 enrichment score, (**E**) L-NMF23 enrichment score in CD49f^+^ATP2B1^+^, CD49f^+^ATP2B1^−^ and CD49f^−^ATP2B1^−^ single cells profiled by scMultiome. Kruskal Wallis test. Box and whiskers. Min to max. (**F**): Unique GSEA terms enriched at a Q value < 0.05 in top driving genes from L-NMF16 signature in CD49f^+^ATP2B1^+^, CD49f^+^ATP2B1^−^ and CD49f^−^ATP2B1^−^ single cells profiled by scMultiome. (Up): GOBP terms, (Bottom): GOCC terms. Dashed red line indicates significance threshold. Bar. (**G**): Predicted terms enriched at a Q value < 0.05 from cis-regulatory region function analysis (GREAT) of the unique peaks in CD49f^+^ATP2B1^+^ single cells profiled by scMultiome. Dashed red line indicates significance threshold. Bar. (**H**): Representative images (bottom) and quantification (top) of confocal analysis of CD49f^+^ATP2B1^+^, CD49f^+^ATP2B1^−^ and CD49f^−^ATP2B1^−^ HSC stained for TFEB and DAPI. Scale bar, 2 μm. n = 3 FL, 352 individual cells/staining; n=3 CB, 377 individual cells/staining, n=3 mPB, 422 individual cells/staining. Scatter dot plot. Mean with SEM. Kruskal-Wallis test. p>0.05 is not shown.

We have previously shown that a TFEB-mediated endolysosomal program governs LT-HSC self-renewal properties. In particular, CB LT-HSC - which display the highest lysosomal content after culture - have superior serial repopulating ability in xenograft assays compared to LT-HSC with lower lysosomal content.^10^ To validate our scMultiome analyses showing enrichment of lysosomal pathways in CD49f^+^ATP2B1^+^ cells, HSC subsets were isolated from CB and lysotracker staining was analyzed by flow cytometry after 16h of culture.^4,24^ CD49f^+^ATP2B1^+^ cells had the highest lysotracker intensity compared to all other fractions (Figure S3H). To determine if the observed enrichment of lysosomal programs in CB CD49f^+^ATP2B1^+^ HSC is conserved across ontogeny, immunofluorescence microscopy was used to quantify TFEB protein in individual cells in HSC subsets from FL, CB and mPB sources.^10^ Total TFEB staining patterns across the three HSC subsets differed across ontogeny, but was consistently lower in CD49f^+^ATP2B1^+^ than CD49f^−^ATP2B1^−^ (Figure 3I). Despite this, the nuclear to non-nuclear fraction of TFEB protein was significantly higher in CD49f^+^ATP2B1^+^ HSC compared to both CD49f^+^ATP2B1^−^ and CD49f^−^ATP2B1^−^ HSC subsets across all human HSC sources tested (Figure 3H). No differences in lysosomal mass as measured by LAMP1 staining were revealed across HSC populations or ontogeny stages, with the exception of a significantly decreased lysosomal mass in mPB-derived CD49f^−^ATP2B1^−^ HSC (Figure S3J). In sum, these data indicate ATP2B1 surface expression marks LT-HSC with enrichment of an TFEB-mediated endolysosomal program.

### CD49f^+^ATP2B1^+^ LT-HSC display superior *in vivo* repopulation capacity and self-renewal ability

To investigate the *in vivo* functionality of HSC subsets marked by surface expression of ATP2B1 and CD49f, equal numbers of CD49f^+^ATP2B1^+^, CD49f^+^ATP2B1^−^ and CD49f^−^ATP2B1^−^ HSC isolated from CB were transplanted into immunodeficient NSG mice. Repopulation was assessed after 4 weeks (4w), a time when LT-HSC are active and proliferating, and at 20 weeks (20w), when repopulation is derived from an LT-HSC compartment that has primarily returned to quiescence (Figure 4A).^4^ Although human leukocyte engraftment from the three HSC subsets was similar at 4w post-transplant (Figures 4B and S4A), the proportion of primitive CD34^+^CD19^−^ was significantly higher in mice engrafted with CD49f^+^ATP2B1^+^ HSC (Figures 4C and S4B). These results are concordant with what was observed in our *in vitro* assays (Figure 2B). By 20w post-transplant, we found that engraftment was significantly higher in recipients transplanted with CD49f^+^ATP2B1^+^ compared to CD49f^+^ATP2B1^−^ HSC. Notably, CD49f^−^ATP2B1^−^ HSC resulted in significantly lower engraftment compared to both CD49f^+^ATP2B1^+^ HSC (5.36-fold decrease) and CD49f^+^ATP2B1^−^ HSC (3.98-fold decrease), indicating that this HSC subset has limited sustained proliferative output capacity (Figures 4D and S4C,D,E). CD49f^+^ATP2B1^+^ cells resulted in grafts with the highest proportion of CD34^+^CD19^−^ cells at 20w (Figures 4E and S4F, G). There was no lineage bias observed within leukocytes (Figure S4H-L), though a higher fraction of GlyA^+^ cells was found in mice engrafted with the CD49f^+^ATP2B1^+^ fraction compared to the other HSC subsets (Figure S4 M,N).

**Figure 4:**
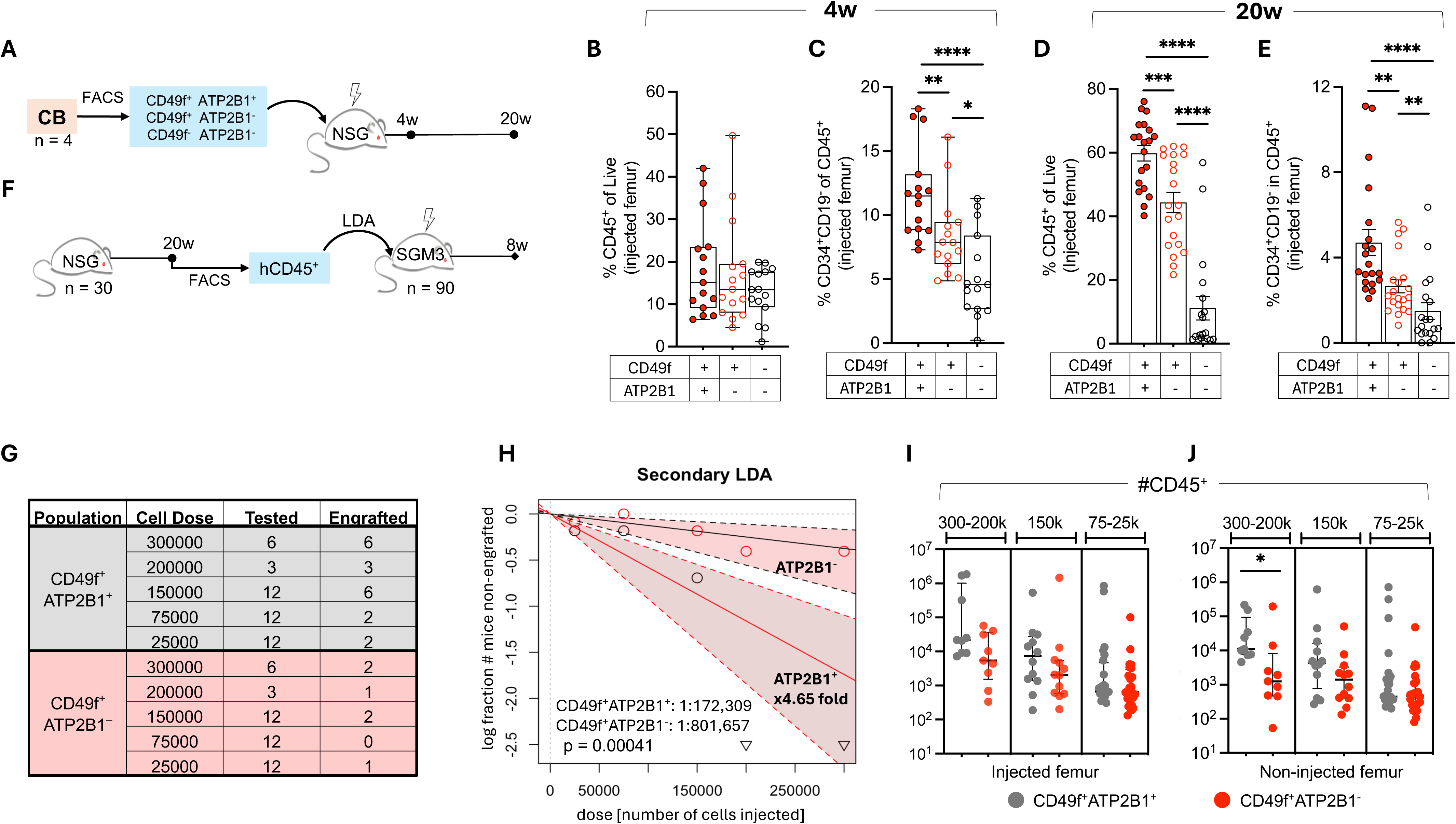
CD49f^+^ATP2B1^+^ LT-HSC display superior *in vivo* repopulation capacity and self-renewal ability. (**A**): Schematic for xenograft assays. CD49f^+^ATP2B1^+^, CD49f^+^ATP2B1^−^ and CD49f^−^ATP2B1^−^ cells were sorted from n=4 CB pools and injected in NSG immunodeficient mice. Hematopoietic organs were analyzed by flow cytometry at 4w and 20w after xenotransplantation. (n=2500 single cells/mouse). (**B-E**): From (**A**): percentage of total CD45^+^ human cells (B,D) in injected femur at 4w (**B**) and 20w (**D**) of engraftment in NSG mice or % of total CD34^+^CD19^−^ cells within CD45^+^ cells (**C,E**) in injected femur at 4w (**C**) and 20w (**E**) of engraftment in NSG mice. Mann-Witney test. Box and whiskers. Min to max show all points. p>0.05 is not shown. (**F**): Schematic for LDA assays. CD45^+^ from bone marrow of NSG mice in Figure 4A at 20w were sorted and injected into secondary NSG-SGM3 recipients at limiting dilution doses. Hematopoietic organs were analyzed by flow cytometry at 8w after xenotransplantation. (n=300000; 200000; 150000; 75000; 25000 cells/mouse). (**G,H**): Graph of HSC frequencies for CD49f^+^ATP2B1^+^ and CD49f^+^ATP2B1^−^ (**H**) and table summarizing results of secondary assays (**G**) with CD45^+^ cells isolated from 30 primary mice and g doses for 8 weeks. n=3 biological replicates. Human CD45^+^ marking of >0.05% in both injected femur and non-injected bones was considered positive for secondary engraftment. p value was by extreme limiting dilution analysis (ELDA).(**I,J**): From (**F**): percentage of total CD45^+^ human cells in injected femur (**I**) and non-injected bones (**J**) at 8w of engraftment in SGM3 mice. Mann-Witney test. Scatter dot plot. Median with interquartile range. p>0.05 is not shown.

To determine if CD49f^+^ATP2B1^+^ cells had superior self-renewal capacity compared to CD49f^+^ATP2B1^−^ HSC, we performed secondary transplantation with limiting dilution assays (LDA) into NSG-SGM3 recipients (Figure 4F,G). Note that serial transplantation of the CD49f^−^ATP2B1^−^ grafts was not attempted here due to low cellularity (Figure S4E). CD49f^+^ATP2B1^+^ grafts resulted in a 4.65-fold increase in HSC frequency compared to those from CD49f^+^ATP2B1^−^ HSC (Figure 4H). All secondary engrafted mice had multilineage reconstitution; yet, the number of engrafted human cells within each lineage tended to be higher in CD49f^+^ATP2B1^+^ over CD49f^+^ATP2B1^−^ across all doses (Figure 4I,J+Figure Se4O-R). Overall, these data validate ATP2B1 as a novel marker for LT-HSC with superior long-term self-renewal.

## Discussion

Here, we demonstrate that ATP2B1 is expressed on the surface of human LT-HSC from across the spectrum of fetal to adult ontogeny. ATP2B1 expression bifurcates the widely used gold standard marker CD49f into two populations. When compared to CD49f^+^ATP2B1^−^ HSC, the CD49f^+^ATP2B1^+^ subset exhibited superior capacity for serial replating of clonogenic progenitors and repopulation and self-renewal in xenografts; HSC frequency was also the highest. ATP2B1 expression and serial clonogenic output was much higher in LT-HSC from human FL followed by a gradual reduction in CB and mPB, a finding consistent with prior studies using less purified CD34 subsets.^19^ For example, ATP2B1 is widely expressed in LT-HSC from FL (mean 46%), but only ∼10% of CB and ∼3-4% of mPB LT-HSC are ATP2B1^+^. Although ATP2B1 expression is heterogeneous across the human lifespan, our data suggests its expression is reflective of functional stemness programs. Thus, interrogation of CD49f^+^ATP2B1^+^ across ontogeny should reveal stemness programs that are common as well as those that change as HSC age.

Recent single-cell transcriptional profiling of CD34^+^ HSPC across the human lifespan have uncovered the dynamic changes associated with human blood progenitors from fetal development through adulthood.^34,35^ These studies reinforced the importance of integrating transcriptional and functional features of HSPC at different developmental stages,^18^ but still did not achieve enough resolution to capture specific facets of LT-HSC biology. To our knowledge, ours is the first study to dissect both the molecular and functional properties of the human CD90^+^ HSC compartment at single-cell resolution. Single-cell transcriptional and epigenetic profiling identified TFEB-mediated control of catabolic processes as one of the molecular mechanisms underpinning the distinct stemness advantage of CD49f^+^ATP2B1^+^-marked HSC across ontogeny. Our findings refine our prior studies that had linked TFEB to CD49f^+^ LT-HSC as compared to CD49f^−^ ST-HSC by showing that TFEB and lysosomal activity preserve quiescence and enhanced self-renewal of CB LT-HSC.^10^ Our new work solidifies the value of quantifying the nuclear versus non-nuclear localization of TFEB - in addition to its total content - as an extra layer of molecular information that helps to identify LT-HSC with superior functionality across the entire spectrum of fetal and adult sources.^10^

By linking surface ATP2B1 expression to endolysosomal LT-HSC programs, our study pinpoints particular pathways for HSC functionality across the human lifespan. Notably, the molecular underpinnings of HSC functional attrition with human aging is unclear, but recent studies point to metabolic and inflammatory stress programs.^25,36–38^ Nonetheless, many functionally young HSC can persist into old age.^39,40^ Autophagy is a catabolic process, centred at the lysosome, directly implicated in HSC dormancy and self-renewal.^41–44^ Our scMultiome and immunostaining analysis suggests that the TFEB-mediated endolysosomal axis is highest in CD49f^+^ATP2B1^+^ cells, nominating ATP2B1-expressing LT-HSC to contain distinct metabolism including enhanced catabolism and autophagy. Although autophagy is decreased overall in murine HSC during aging, a subset of aged HSC activates autophagy to counteract aging- and inflammation-associated metabolic impairment.^39^ We hypothesize that these represent the functional ATP2B1^+^CD49f^+^ HSC we have identified in humans, but also potentially pinpoints why we find their number decreased by ∼3-fold in mPB as compared to CB. When robust low-input to single-cell metabolomics profiling is available to directly profile metabolic processes in ATP2B1^+^ and ATP2B1^−^ LT-HSC, such future studies should shed considerable light to human HSC heterogeneity. Overall, our findings serve as a foundation for future studies aimed at examining the molecular and metabolic programs governing human LT-HSC across ontogeny.

### Limitations of the Study

In our study, we demonstrate that ATP2B1 surface expression allows purification of functionally distinct human LT-HSC across ontogeny. Because immunodeficient mice, FL, and mPB are limited resources, we relied on CB as a representative source for our gold standard xenotransplantation assays as well as scMultiomics.

We have not determined if the expression of ATP2B1 is sufficient to endow stemness properties onto human HSPC; genetic studies for overexpressing ATP2B1 in primary cells are challenging owing to the larger size of the ATP2B1 transgene (>3.5kb). ATP2B1 also has 17 transcript variants, and we have not been able to isolate which are expressed in HSC or important for the phenotypes observed in this study. Similarly, any genetic perturbation of ATP2B1 would require *in vitro* culture which has been shown to trigger immune-mediated inflammatory pathways and genetic aberrations not observed at homeostasis.^45^ To overcome some of these challenges, we functionally validated the role of ATP2B1 in our assays using the selective small molecule inhibitor PI-8 in unperturbed settings. This avoided *in vitro* pre-culture, electroporation, and/or CRISPR/Cas9-mediated manipulation of the genome.

## Lead contact

Further information and requests for resources and reagents should be directed to and will be fulfilled by the lead contact, Dr. Stephanie Z. Xie.

## Materials availability

All unique materials generated in this study are available from the lead contact upon reasonable request with a completed materials transfer agreement.

## Acknowledgments

We thank the obstetrics unit of Trillium Health, William Osler and Credit Valley Hospitals for CB; L.Q., K.K. and A.M. for help with intra-femoral injections; K.K. for help with immunofluorescence analysis, O.G. for help with the single-cell stroma assays, the Leukemia Tissue Bank at Princess Margaret Cancer Centre for sample coordination, the UHN-Sickkids Flow cytometry facility for cell sorting and the Princess Margaret Genomic Centre for 10x Multiome library prep and sequencing.

## Author contributions

Conceptualization of a surface LT-HSC marker study: M.S.N., A.G.X.Z., and S.Z.X.; Conceptualization of an human ontogeny LT-HSC comparative study: S.Z.X.; Methodology, A.V., M.S.N, and S.Z.X.; Investigation, A.V., M.S.N, I.D.B., H.K., M.Z. and S.Z.X.; Formal analysis, A.V., S.S., A.G.X.Z., and A.M.; Visualization, A.V., I.D.B., M.S.N., S.S. and S.Z.X.; Supervision, J.E.D and S.Z.X.; Funding acquisition, J.E.D., and S.Z.X.; Writing - Original Draft, A.V. and S.Z.X; Writing - Review & Editing, all authors.

## Materials and Methods

### Cord Blood (CB), Fetal Liver (FL) and Mobilized Peripheral Blood (mPB) samples

All human CB and mPB samples were obtained with informed consent according to procedures approved by the institutional review boards of the University Health Network, and where applicable, Trillium Health Centre, Brampton Civic Hospital and Credit Valley Hospital, Ontario, Canada. Mononuclear cell (MNC) isolation and subsequent CD34^+^enrichment of CB was conducted as previously described.^25^ Human mPB samples (female and male, aged 22-65 years) from G-CSF-treated healthy donors were obtained via the Leukemia Tissue Bank at Princess Margaret Cancer Centre according to standard procedures. Following MNC isolation, CD34^+^ cells were enriched as for CB. FL were obtained, processed, and CD34^+^ enriched as previously described.^46^ Cells were cryopreserved at −150C.

### Flow cytometry and cell sorting

FL, CB, and mPB CD34^+^ enriched samples were thawed by dropwise addition of X-VIVO 10 + 50% fetal bovine serum supplemented with DNase I (Roche; 100 μg/ml final concentration) and resuspended at a density of less than or equal to 10 × 10^6^ cells/mL in PBS with 5%v/v FBS. Cells were stained for 15 minutes at RT with the following mouse anti-human antibodies: anti-CD19-BV711, anti-CD34-APC-Cy7, anti-CD38-PE-Cy7, anti-CD45RA-FITC, and anti-CD90-APC. Additionally, rat anti-human CD49f-PE-Cy5 was used, along with mouse anti-human ATP2B1-MaxLight550. After washing, cells were resuspended at 0.5–10 × 10^6^ cells/mL in PBS with 2% FBS and 0.1 μg/mL propidium iodide (PI). Cells were sorted on BD FACSAria Fusion or BD Symphony S6 instruments. LT-HSC and ST-HSC were sorted essentially as previously described.^4,5,10^ ATP2B1-MaxLight550 FMO controls were used to establish gates for CD49f^+^ATP2B1^+^, CD49f^+^ATP2B1^−^ and CD49f^−^ATP2B1^−^in each sample.

### Colony forming cell assay

250 CD49f^+^ATP2B1^+^, CD49f^+^ATP2B1^−^ and CD49f^−^ATP2B1^−^ HSC from FL and CB and 1500 CD49f^+^ATP2B1^+^, CD49f^+^ATP2B1^−^ and CD49f^−^ATP2B1^−^ HSC from mPB were sorted into 2.5mL of Methocult H4034 optimum methylcellulose media (cytokines: SCF, GM-CSF,IL-3, G-CSF, EPO, STEMCELL Technologies) supplemented with FLT3 ligand (10 ng/mL) and IL-6 (10 ng/mL) for the primary assay. After thorough resuspension, 1mL of methocult was plated into 35 mm dishes in duplicate (100 cells/dish). Colonies were allowed to differentiate for 12 days and were counted and morphologically assessed. On day 14 after plating, cells were harvested from the methocult, washed and resuspended in 1mL of PBS. For serial replating of the primary colony assay, 1%v/v for FL and CB, or 3%v/v for mPB of cell suspension was seeded into 2.5mL of Methocult and plated as above. Colonies were allowed to grow for 12 additional days and were then counted and morphologically assessed. Flow cytometry was also carried out on harvested primary and secondary colonies at day 14 and 28, respectively from initial plating. CD34^+^ cells were quantified by staining the cells for 15 minutes at RT with mouse anti-human CD34-APC-Cy7. Additionally, the following mouse anti-human antibodies were included for lineage validation only: anti-CD45RA-FITC, anti-CD14-BV605, anti-GlyA-PE, anti-CD15-PE-Cy5, anti-CD66b-AlexaFluor647, anti-CD45-AlexaFluor700 and anti-CD33-BV786. Cells were washed, stained with PI and analyzed on a BD Symphony A1 instrument.

### Single-cell differentiation assays on MS5 stroma

Single-cell *in vitro* differentiation assays were conducted as previously described.^8^ For index sorting of single HSC, FL, CB or mPB CD34-enriched cells were thawed and stained as above. CD19^−^CD34^+^CD38^−^CD45RA^−^CD90^+^ single cells were sorted onto established MS5 stromal layers using a BD FACSAria Fusion. Fluorescence intensities for channels reflecting CD49f and ATP2B1 expression were indexed. Single cells were differentiated for 18d in erythromyeloid (EM) medium as previously described.^8^ Cloning efficiency was determined by counting the number of wells with growth of cells on day 17. The following antibody panel was used for flow cytometry analysis on day 18: GlyA-PE, CD71-FITC, CD34-APC-Cy7, CD33-BV786, CD66b-AF647, CD14-PECy7, CD56-BV605, CD41-PECy5 and CD45-AF700. Sytox blue (Thermo Fisher) was used as a viability stain. Data was acquired using a BD Symphony A1 and analyzed on FlowJo v10 to determine clonal lineage output, which was aligned with indexed single-cell fluorescence intensity for ATP2B1 at initial single-cell deposition.

### Immunofluorescence microscopy

CD49f^+^ATP2B1^+^, CD49f^+^ATP2B1^−^ and CD49f^−^ATP2B1^−^ cells were prepared for immunostaining as described previously.^10^ The following primary antibodies were used: rabbit anti-ATP2B1, AF488 anti-LAMP1, and mouse anti-TFEB. The following secondary antibodies were used: AF568 donkey anti-rabbit, AF568 goat anti-mouse. Images of individual cells were captured with a Zeiss LSM700 confocal microscope (oil, ×63/1.4 NA, Zen 2012) or a Leica LP8 confocal microscope (HC PL APO 63x/1.40 NA Oil immersion CS2), and images were processed and analyzed with ImageJ/Fiji as previously described.^8^

### Mice

Animal experiments were done in accordance with institutional guidelines approved by the UHN Animal care, and we complied with all relevant ethical regulations for animal testing and research. NOD.Cg-*Prkdc^scid^Il2rg^tm1Wjl^*/SzJ(NSG), and NOD.Cg-*Prkdc^scid^Il2rg^tm1Wjl^*Tg(CMV-IL3,CSF2,KITLG)1Eav /MloySzJ (NSG-SGM3) mice were housed in a controlled environment with a 12-h:12-h light:dark cycle including a 30-min transition, a room temperature (RT) of 21–23 °C and a humidity of 30–60%, and had *ad libitum* access to dry laboratory food and water at the animal facility (ARC) at Princess Margaret Cancer Centre. Experimental mice were housed in a room designated only for immunocompromised mice with individually ventilated racks equipped with complete sterile micro-isolator caging (IVC), on corn-cob bedding and supplied with environmental enrichment in the form of a red house/tube and a cotton nestlet. Cages were changed every <7 d under a biological safety cabinet. Health status was monitored using a combination of soiled bedding sentinels and environmental monitoring.

### Xenotransplantation

Age- and sex-matched 8- to 12-week-old male and female recipient NSG mice were sublethally irradiated with 225 cGy of gamma radiation (GammaCell40) 1 day before engraftment. HSC (CD19^−^CD34^+^CD38^−^CD45RA^−^CD90^+^) were fractionated into the indicated populations by CD49f and ATP2B1 expression using FACS. Sorted cells were injected by intrafemoral injection. Mice were euthanized at the indicated time points post-engraftment, and the injected femur and other long bones (non-injected femur and tibiae) were flushed separately in Iscove’s modified Dulbecco’s medium (IMDM) supplemented with 5% FBS. Typically, 10% of the total collected cells were stained for flow cytometry. Spleens were also isolated, macerated in IMDM+5%FBS, filtered (30uM), and 6.5% of total collected cells were stained for flow cytometry. The following antibodies were used to evaluate engraftment: CD14-PECy5, GlyA-PE, CD33-BV786, CD45-AF700, CD66b-AF647, CD34-APCCy7, CD19-PE, CD45-V500, CD56-BV605, CD3-FITC. Cells were stained and washed as above, with Sytox Blue (Thermo Fisher) used to gauge viability and 75% of the stained cells were analyzed on a BD Symphony A1. The remainder of isolated cells were pooled by condition and cryopreserved at -80C.

### Secondary Transplantation

Cryopreserved bone marrow from primary transplanted mice as above were thawed as described. Mouse cell depletion was carried out using a kit (Miltenyi) per the manufacturer’s protocol. Human leukocytes were further purified by FACS using hCD45 as a marker. For secondary transplantation at limiting dilution doses, human CD45^+^ cells were injected by intrafemoral injection at the indicated doses into irradiated (225cGy) male or female 8-12w-old NSG-SGM3 mice. Engraftment was assessed at 8 weeks after transplant. Injected and contralateral femurs were isolated, flushed with IMDM+5%FBS and stained separately with the antibody panel indicated above. A mouse was considered engrafted if the percentage of human CD45 cells in the live gate was >0.05 in both femurs. HSC frequency was estimated using ELDA (http://bioinf.wehi.edu.au/software/elda/).

### Statistical analysis

GraphPad Prism (version 10.2.3) was used for all statistical analyses, unless otherwise noted. Statistical significance (\**p* < 0.05, \*\**p* < 0.01, \*\*\**p* < 0.001 and \*\*\**p* < 0.0001) was determined using the statistical methods reported in the respective figure legends or otherwise noted. Typically, if normal-distribution assumptions were not valid, Mann–Whitney *U*-tests (two-tailed) or Kruskal–Wallis tests were done for single or multiple comparisons, respectively. Kolmogorov–Smirnov tests were done to test for normality, if the sample size allowed. One-way ANOVAs were used to compare means among three or more independent groups when normality was assumed. Multiple testing corrections and post-hoc tests were completed to compare all pairs of treatment groups when the overall *p*-value < 0.05. Unless otherwise noted, *p*-value > 0.05 is not shown.

### LysoTracker analysis

After sorting, HSC populations as CD49f^+^ATP2B1^+^, CD49f^+^ATP2B1^−^, CD49f^−^ATP2B1^−^, cells were resuspended in X-VIVO 10 medium supplemented with 1% BSA (Roche), SCF (100 ng/ml), FLT3L (100 ng/ml), TPO (50 ng/ml) and IL-7 (10 ng/ml), 1x L-glutamine (Thermo Fisher), 1x penicillinstreptomycin (Thermo Fisher), and cultured for 30 min at 37**°**C with 75nM LysoTracker. After staining, cells were washed once and resuspended in PBS+2% FBS and analyzed on BD SymphonyA1. FlowJo v10 was used for analysis.

### Single-cell (sc)Multiome (RNA and ATAC) sorting and library preparation

For scMultiome profiling, cryopreserved CB from multiple donors was pooled and stained for FACS as described above and sorted for CD49f^+^ATP2B1^+^, CD49f^+^ATP2B1^−^, CD49f^−^ATP2B1^−^ HSC. The three populations were processed for nuclei isolation and library construction by the Princess Margaret Genome Centre for downstream 10x Genomics Chromium scMultiome scRNA + scATAC sequencing using their standard protocol.

### scMultiome preprocessing

scMultiome data generated from previous studies and shown in Figure S1A were processed as described therein.^25^ scMultiome data generated in this study were aligned to GRCh38 using CellRanger-ARC (v2.0.2), which was subsequently used to aggregate samples and normalize read counts to a uniform median depth. For each sample, doublets were identified using scDblFinder (v1.12.0)^47^ and AMULET,^48^ with default parameters. Doublets identified by either scDblFinder or AMULET were removed from downstream analysis. Signac (v1.11.0)^49^ and Seurat (v4.3.0)^50^ were used to read in data and perform QC for ATAC- and RNA-seq. The following QC metrics were used: percentage of mitochondrial genes (pct_mito), percentage of ribosomal genes (pct_ribo) nucleosome banding pattern (nucleosome_signal), ATAC transcriptional start site enrichment (TSS_enrichment), minimum number of unique RNA transcripts detected (min_nCountRNA), maximum number of unique RNA transcripts detected (max_nCountRNA), minimum number of fragments in peaks (min_nCountATAC) and maximum number of fragments in peaks (max_nCountATAC). Thresholds for QC filtering were adapted to each dataset based on the distribution of QC metrics. For CD49f^+^ATP2B1^+^, filtering thresholds were set at: min_nCountRNA>2500, max_nCountRNA<20000, min_nCountATAC>6500, maxnCountATAC<100000, nucleosome_signal<1.1, TSS_enrichment>3, pct_mito<20, pct_ribo>5. For CD49f^+^ATP2B1^−^, filtering thresholds were set at: min_nCountRNA>2500, max_nCountRNA<19000, min_nCountATAC>6500, maxnCountATAC<95000, nucleosome_signal<1.1, TSS_enrichment>3, pct_mito<20, pct_ribo>5. For CD49f^−^ATP2B1^−^, filtering thresholds were set at: min_nCountRNA>2500, max_nCountRNA<15000, min_nCountATAC>6500, maxnCountATAC<95000, nucleosome_signal<1.4, TSS_enrichment>3, pct_mito<20, pct_ribo>8.

The RNA feature count matrix was corrected for ambient RNA contamination using SoupX (v1.6.2)^51^ with default parameters. Donor sex and identity from the pooled scMultiome was inferred as previously described.^25^

### scMultiome Processing and Classification

After initial quality control, the pooled scMultiome was normalized by SCTransform (v0.3.5), followed by variable feature selection, scaling, and PCA reduction. Batch correction with harmony (v1.2.0)^52^ was performed based on the sample identity. Data were processed using the top 50 harmony-corrected principal components, which were used to construct a neighborhood graph using the top 30 neighbors for each cell and perform UMAP reduction. FindMarkers function from Seurat (v4.3.0)^32^ was used to determine the top marker genes for each sample using standard parameters. For chromatin analyses, peaks were called using MACS2^53^ from fragment files corresponding to high quality cells after QC filtering, and bedtools was used to identify the maximum fold enrichment observed at each peak in the catalog in each population. Peaks within non-standard chromosomes and genomic blacklisted regions were removed from downstream analyses. From the resulting peak matrix, dimensionality reduction with latent semantic indexing (LSI) was performed following the Signac pipeline and LSI components were batch corrected by sample identity using harmony. Harmony-corrected LSI components 2 to 40 were used to construct a neighborhood graph using the top 30 neighbors for each cell and perform UMAP reduction. For integrative analysis, weighted nearest neighbors (WNN) integration^32^ was performed using harmony-corrected RNA PCA components 1:50 and harmony-corrected ATAC LSI components 2:40, considering the top 30 neighbors for each cell. UMAP reduction was performed with a min.dist=0.2 along the WNN neighborhood graph.^32^ Homer was used to identify enriched motifs within the unique peak sites for each sample, using default parameters, and the catalog of all called peaks as a background. GREAT (v.4.0.4, http://great.stanford.edu/public/html/), was used to predict the functions of cis-regulatory regions using the unique peaks per sample as Test regions, the catalogue of all peaks as Background and the “basal plus extension” option with default parameters.

### RNA and ATAC signature scoring

Geneset scoring of previously published gene expression signatures was performed as previously described.^25^ Enrichment of chromatin regions in scATAC-seq data was evaluated through chromVAR (v1.20.2).^31^ In all cases, standardization was performed across cells prior to plotting for ease of visualization. ChromVAR ZScores enrichments higher than 1.96 was used to classify cells positive for a given signature providing no others were enriched (i.e. LT/HSPC or Act/HSPC).

Linked non-negative matrix factorization (L-NMF)^33^ was run with default parameters indicating sample of origin as variable for batch correction. The top driving genes of each L-NMF signature was determined by identifying which genes were higher than the inflection point on an elbow plot of ranked feature weight.

### Gene set enrichment analysis

Gene set enrichment analysis (GSEA) was performed on the top genes driving each L-NMF signature using gProfiler (https://www.google.com/search?client=safari&rls=en&q=gprofiler&ie=UTF-8&oe=UTF-8) with default parameters. GO molecular function, GO cellular component and GO biological processes were used as score datasets on the top genes from each LNMF signature. Pathways with Q value<0.05 were selected as significant and used for subsequent analysis. Unique pathways representative of each L-NMF signature were selected with Venny (v2.1.0, https://bioinfogp.cnb.csic.es/tools/venny/) and used for representation.

### Bulk RNA-seq library preparation and processing

Bulk RNA-seq data shown in Figure S1A were obtained and processed as previously described.^10^

---

**Figure S1:**
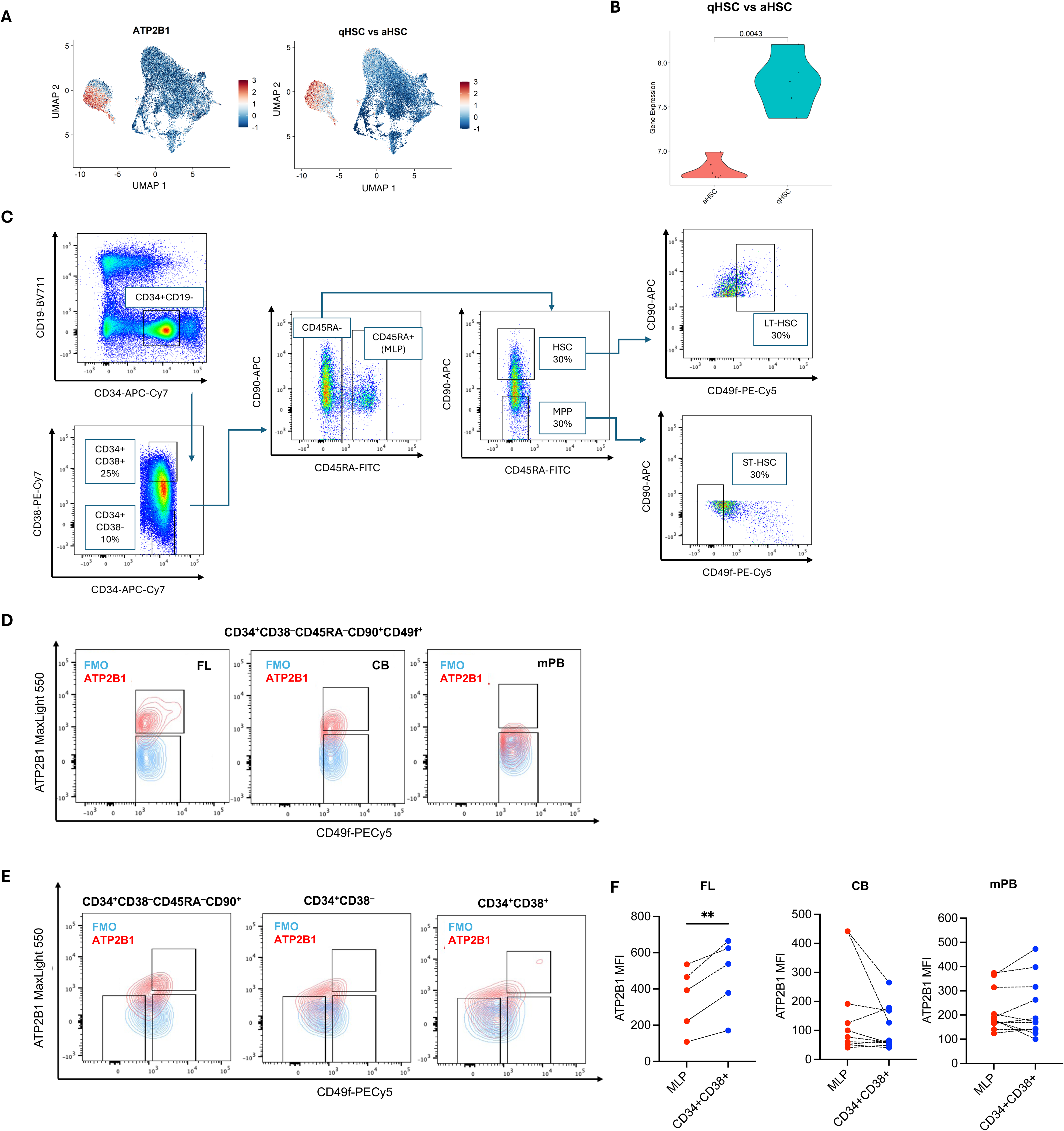
(**A**): Integrated WNN (RNA+ATAC) UMAP embedding showing (left) normalized *ATP2B1* expression, (right) Normalized enrichment score (AUCell) of the qHSC vs aHSC in CB-CD34^+^CD38^−^CD45RA^−^ HSPC profiled by scMultiome at 20w after xenotransplantation.^25^ CB = single cells. Colors indicate the degree of enrichment for each signature. (**B**): Normalized *ATP2B1* gene expression in qHSC and aHSC from CB HSPC populations profiled by bulk RNAseq.^10^ Mann-Whitney test. Violin plot. (**C**): Flow cytometry and sorting scheme for the indicated populations from CD34-column enriched human FL, CB and mPB. (**D**): Representative flow cytometric plots of ATP2B1^+^ and ATP2B1^−^ cells within human CD34^+^CD38^−^CD45RA^−^CD90^+^CD49f^+^ HSC in FL, CB and mPB. Light blue: FMO: Fluorescent Minus One; red: ATP2B1 stained cells. (**E**): Representative flow cytometric plots of ATP2B1^+/–^ and CD49f^+/–^ cells within the indicated human HSPC subpopulations in FL, CB and mPB. Light blue: FMO: Fluorescent Minus One; red: ATP2B1 stained cells. (**F**): ATP2B1 MFI in MLP and CD34^+^CD38^+^cells analyzed after thawing by flow cytometry. n=4 FL, n=10 CB, n=10 mPB. Ratio paired t test.p>0.05 is not shown. Before-after. Symbols & lines.

**Figure S2.**
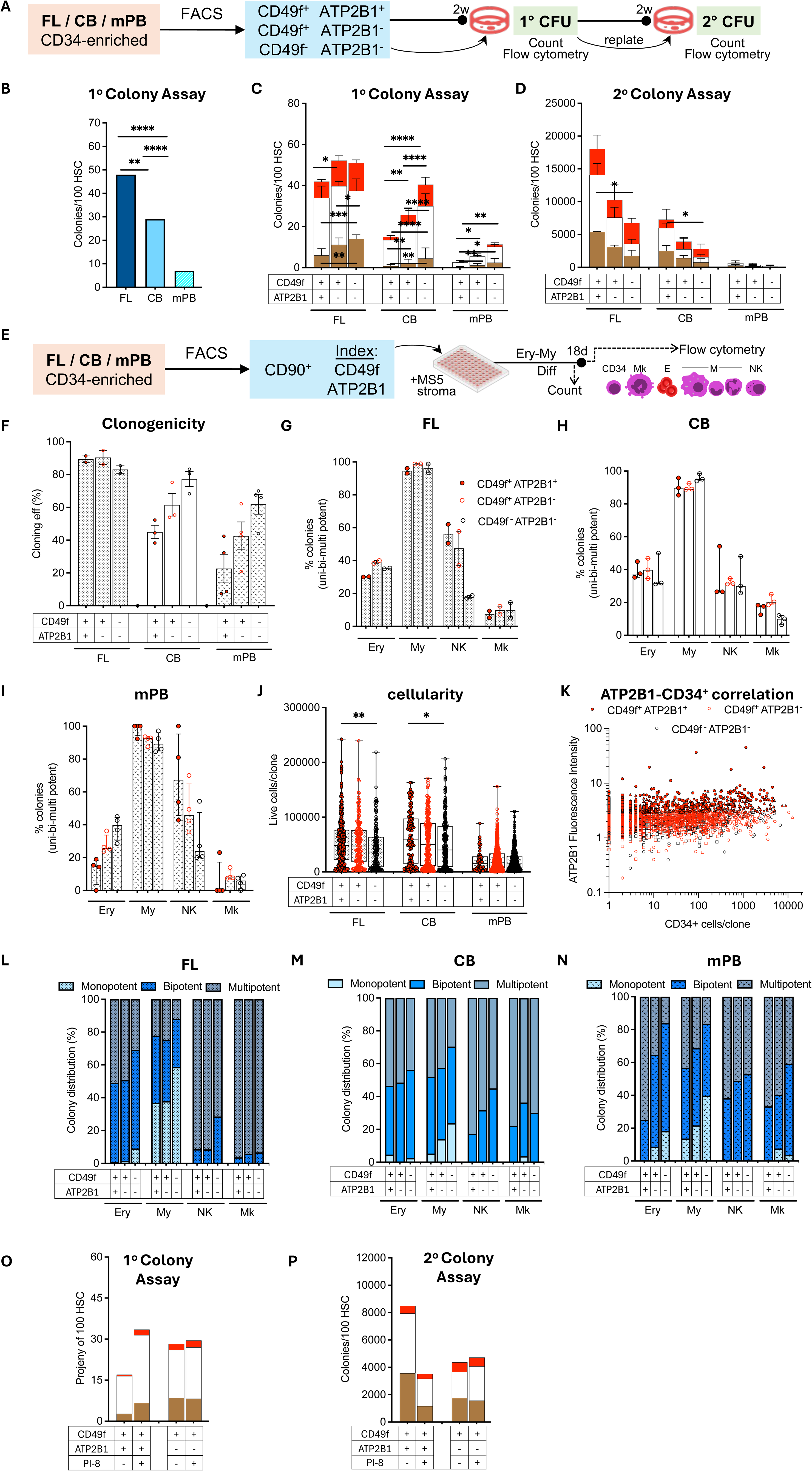
(**A**): Experimental scheme. The indicated HSC subsets were sorted, grown for 14 days in a methylcellulose-based media, counted, stained, assessed by flow cytometry. Serial replating of primary colonies was performed at day 28. n=5 FL, n=14 CB, n=10 mPB. (**B**): Average percentage of cloned and non-cloned colonies from CD49f^+^ATP2B1^+^, CD49f^+^ATP2B1^−^ and CD49f^−^ATP2B1^−^ HSC subsets shown in Figure 2A. n=4 FL, n=10 CB, n=10 mPB. Fisher’s exact test. Bar. (**C,D**): CD49f^+^ATP2B1^+^, CD49f^+^ATP2B1^−^, CD49f^−^ATP2B1^−^ and ST-HSC clonogenic assays showing the lineage output of (**C**) primary colonies from Figure 2A; (**D**) secondary colonies from Figure 2C. Two-way ANOVA statistics. p>0.05 is not shown. Bar. Median with interquartile range. (**E**): Experimental scheme. HSC were index sorted, cultured for 18 days on a stroma-supported erythroid myeloid media, stained and accessed by flow cytometry. n=1280 single FL HSC, n=1858 single CB HSC, n=5036 single mPB HSC. (**F**): Percentage of single cells that grew into a colony in the single cell in vitro differentiation assays from FL (n=2), CB (n=3) and mPB (n=4) experiments with independent single donors (FL and mPB) or donor pools (CB). Scatter dot plot. Mean with SEM. (**G-I**): Percentage of single cells that grew into a colony containing Erythroid (Ery), Myeloid (My), NK (CD56^+^) or Mk (Megakariocitic) cells in the single cell in vitro differentiation assays from (**G**) FL (n=2), (**H**) CB (n=3) and (**I**) mPB (n=4) experiments with independent single donors (FL and mPB) or donor pools (CB). Bar. Median with interquartile range. (**J**): Proliferative potential per colony in the single cell in vitro differentiation assays from FL (n=2), CB (n=3) and mPB (n=4) experiments with independent single donors (FL and mPB) or donor pools (CB). Kruskal Wallis test. p>0.05 is not shown. Box and whiskers. Min to max show all points. (**K**): ATP2B1 MFI of index sorted individual Fl, CB and mPB HSC that grew into single colonies containing CD34^+^ cells at the indicated output in the single cell in vitro differentiation assays from FL (n=2), CB (n=3) and mPB (n=4) experiments with independent single donors (FL and mPB) or donor pools (CB). (**L-N**): Percentage of single cells per lineage that grew into single colonies of only that lineage (Monopotent), one additional (Bipotent), two or more additional (Multipotent) lineages in the single cell in vitro differentiation assays from (**L**) FL (n=2), (**M**) CB (n=3) and (**N**) mPB (n=4) experiments with independent single donors (FL and mPB) or donor pools (CB). Bar. (**O,P**): CD49f+ATP2B1+ and CD49f+ATP2B1- clonogenic assays showing the lineage output of (**O**) primary colonies from Figure 2H; (**P**) secondary colonies from Figure 2I. Two-way ANOVA. p>0.05 is not shown. Bar. Median with interquartile range.

**Figure S3.**
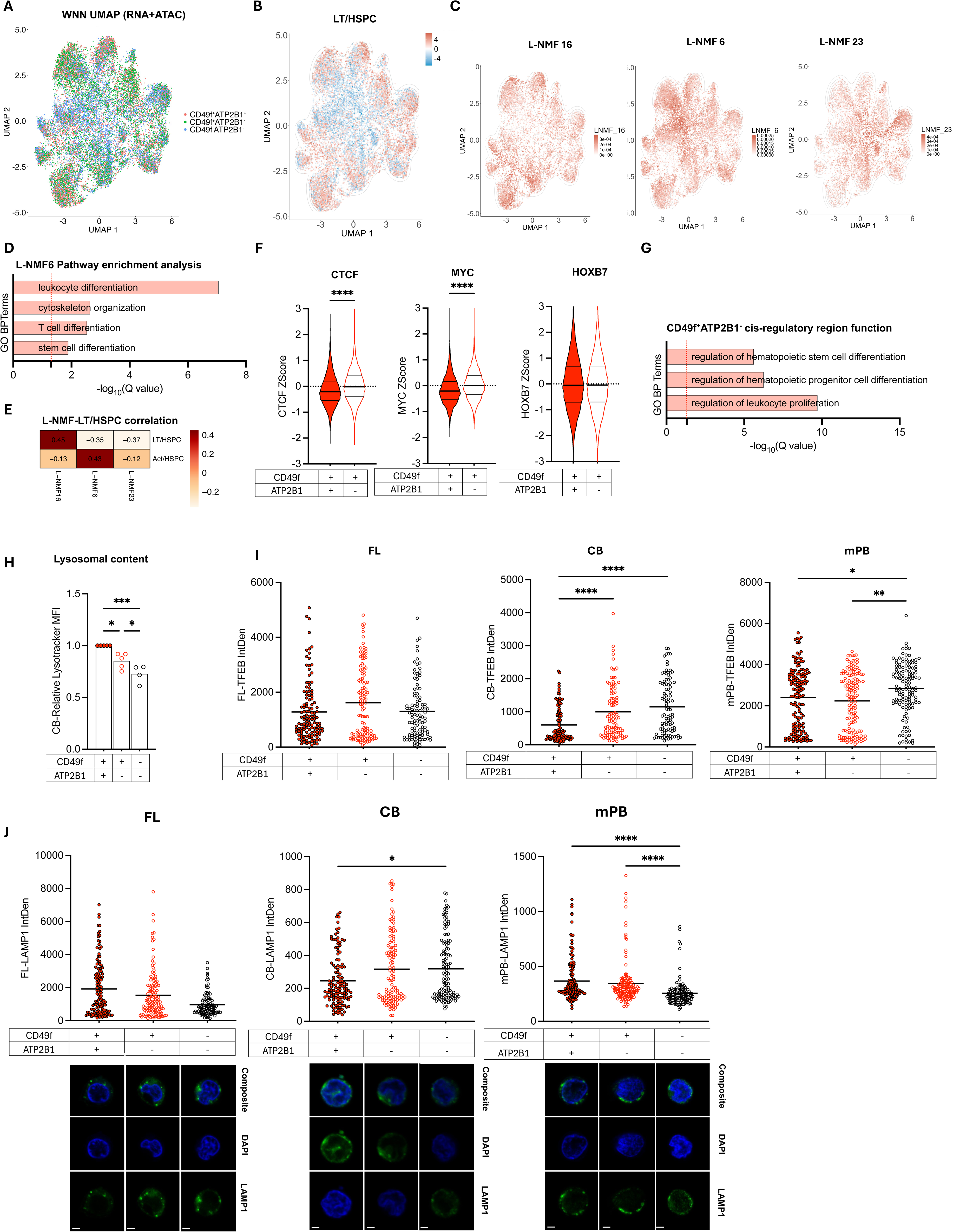
(**A**): Integrated WNN (RNA+ATAC) UMAP embedding showing projection of CD49f^+^ATP2B1^+^, CD49f^+^ATP2B1^−^ and CD49f^−^ATP2B1^−^ single cells profiled by scMultiome. Different colors indicate each HSC subset. (**B,C**): Integrated WNN (RNA+ATAC) UMAP embedding showing (B) enrichment of the LT/HSPC epigenetic signature as in Figure 3A, (C) enrichment of (left) L-NMF16, centre (L-NMF6) and (right) L-NMF23 in CD49f^+^ATP2B1^+^, CD49f^+^ATP2B1^−^ and CD49f^−^ATP2B1^−^ single cells profiled by scMultiome. Colors indicate the degree of enrichment for each signature. (**D**): Unique GSEA terms enriched at a Q value < 0.05 in top driving genes from L-NMF6 signature in CD49f^+^ATP2B1^+^, CD49f^+^ATP2B1^−^ and CD49f^−^ATP2B1^−^ single cells profiled by scMultiome. Dashed red line indicates significance threshold. Bar. (**E**): Heatmap of the strength of Pearson’s correlation for each L-NMF-derived signature (rows) in each chromatin signature (columns,) from lowest (blue) to highest (red) values for LT/HSPC and Act/HSPC unique chromatin peaks. (**F**): *Z* scores for enrichment of CTCF, MYC, and HOXB7 binding sites identified by the ReMap project in CD49f^+^ATP2B1^+^ and CD49f^+^ATP2B1^−^ single cells profiled by scMultiome. Mann Whitney test. Violin plot (truncated). p>0.05 is not shown. (**G**): Predicted terms enriched at a Q value < 0.05 from cis-regulatory region function analysis (GREAT) of the unique peaks in CD49f^+^ATP2B1^−^ single cells profiled by scMultiome. Dashed red line indicates significance threshold. Bar. (**H**): LysoTracker analyzed by flow cytometry in CD49f^+^ATP2B1^+^, CD49f^+^ATP2B1^−^ and CD49f^−^ATP2B1^−^ HSC. n = 4-5 CB. One Way Anova test. Scatter dot plot. Mean with SEM. (**I**): Quantification of confocal analysis of CD49f^+^ATP2B1^+^, CD49f^+^ATP2B1^−^ and CD49f^−^ATP2B1^−^ HSC stained for TFEB and DAPI as for Figure 2H. Scale bar, 2 μm. n = 3 FL, 352 individual cells/staining; n=3 CB, 377 individual cells/staining, n=3 mPB, 422 individual cells/staining. Scatter dot plot. Mean with SEM. Kruskal-Wallis test. p>0.05 is not shown. (**J**): Representative images (bottom) and quantification (top) of confocal analysis of CD49f^+^ATP2B1^+^, CD49f^+^ATP2B1^−^ and CD49f^−^ATP2B1^−^ HSC stained for LAMP1 and DAPI. Scale bar, 2 μm. n = 3 FL, 352 individual cells/staining; n=3 CB, 377 individual cells/staining, n=3 mPB, 422 individual cells/staining. Scatter dot plot. Mean with SEM. Kruskal-Wallis test. p>0.05 is not shown.

**Figure S4.**
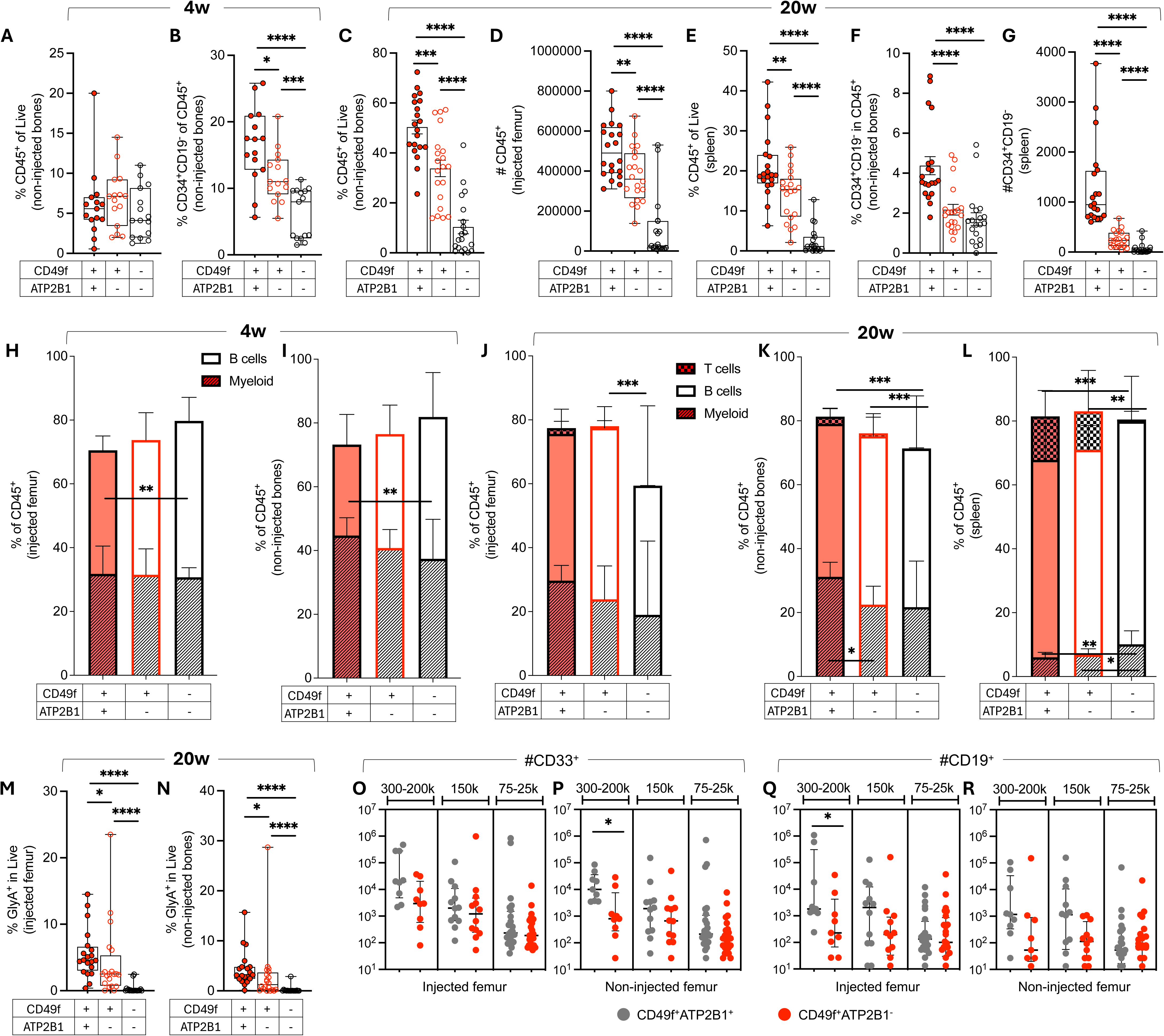
(**A,B,C,F**): From Figure 4A: percentage of total CD45^+^ human cells (**A,C**) in non-injected bones at 4w (**B**) and 20w (**D**) of engraftment in NSG mice or % of total CD34^+^CD19^−^ cells within CD45^+^ cells (**C,E**) in non-injected bones (**B,F**) at 4w (**B**) and 20w (**F**) of engraftment in NSG mice. Mann-Witney test. Box and whiskers. Min to max show all points. p>0.05 is not shown. (**D**): From Figure 4A: number of total CD45^+^ human cells in injected femur at 20w of engraftment in NSG mice. Mann-Witney test. Box and whiskers. Min to max show all points. p>0.05 is not shown. (**E,G**): From Figure 4A: percentage of total CD45^+^ human cells (**E**) in spleen at 20w of engraftment in NSG mice or number of total CD34^+^CD19^−^ cells (**G**) in spleen at 20w of engraftment in NSG mice. Mann-Witney test. Box and whiskers. Min to max show all points. p>0.05 is not shown. (**H-L**): Lineage composition by myeloid (CD33^+^), B (CD19^+^), and T (CD3^+^) cells (**H-L**) in injected, non-injected bones, and spleen from NSG xenografts in Figure 4A at 4w (**H,I**) or 20w (**J-L**) after xenotransplantation. n=4 CB. Mann-Witney test. Bar. Median with interquartile range. p>0.05 is not shown. (**M,N**): Proportion of GlyA^+^ cells in Live in injected (**M**) and non-injected bones (**N**) from NSG xenografts in Figure 4A at 20w after xenotransplantation. n=4 CB. Mann-Witney test. Box and whiskers. Min to max show all points. p>0.05 is not shown. (**O-R**): From Figure 4F: percentage of total CD33^+^ cells (**O,P**) in injected femur (**O**) and non-injected bones (**P**) at 8w of engraftment in SGM3 mice, or percentage of total CD19^+^ cells (**Q,R**) in injected femur (**Q**) and non-injected bones (**R**) at 8w of engraftment in SGM3 mice. Mann-Witney test. Scatter dot plot. Median with interquartile range. p>0.05 is not shown.

